# Fusion Neural Network (FusNet) for predicting protein-mediated loops

**DOI:** 10.1101/2023.06.24.546360

**Authors:** Li Tang, Wenjie Huang, Matthew C. Hill, Patrick T. Ellinor, Min Li

## Abstract

The organization of the three-dimensional (3D) genome is a complex, and requires a plethora of proteins to ensure the proper formation and regulation of chromatin loops as well as higher order structures. Studying protein-mediated loop regulation can help unravel the intricate interplay between these loops and their crucial roles in modulating gene expression across different cellular contexts. However, current targeted chromatin conformation capture experiments face limitations in capturing protein-mediated loops across various cell types, and existing computational methods fail to predict diverse protein-mediated loops. To address these issues, we propose a fusion neural network (FusNet) designed for predicting protein-mediated loops. FusNet leverages genome sequence information, open chromatin, and ChIP-seq data to efficiently represent and analyze the positions of loop anchors. To extract informative features and reduce the complexity of FusNet, we constructed a convolutional neural network, which compresses the dimensionality of the features while also preserving the most significant ones. To enhance the accuracy and generalization capacity of FusNet, we built a fusion layer by stacking the prediction of fundamental models with a meta-model. FusNet demonstrated its effectiveness in predicting protein-mediated loops, exhibiting high consistency with Hi-C data. Moreover, we find that the loops output from FusNet are highly associated with regulatory functions. Through association analysis with genetic risk variants, FusNet further revealed its potential for unraveling disease-related mechanisms. In conclusion, our study offers a novel computational approach for predicting various protein-mediated chromatin loops, which could substantially enhance research on the functional significance of protein-mediated loop structures in diverse cellular contexts.

**Significance Statement:** The intricate spatial organization of the three-dimensional (3D) genome involves functional proteins critically contributing to chromatin loop formation and regulation. Understanding these protein-mediated loops is vital for elucidating their influence on 3D genome architecture and gene regulation across different cellular types and disease-related contexts. In this study, we propose a Fusion Neural Network (FusNet) for predicting protein-mediated loops. FusNet can concurrently capture and analyze multiple protein-mediated loops in various cell types to advance our understanding of the multitude of protein-mediated loop structures and their functional significance. Importantly, through association analysis with risk variants, FusNet manifests potential in revealing disease-related mechanisms.

## Introduction

In eukaryotic organisms, chromatin typically folds into a three-dimensional structure, which influences gene regulation, expression, cellular function, and ultimately impacts a multitude of biological processes and developmental events (1, 2). Recent advancements in Chromosome Conformation Capture (3C)-based methodologies have facilitated the elucidation of the 3D genomic architecture (3–6). The emergence of targeted chromatin conformation capture techniques enables the detection of protein-specific chromatin interactions. ChIA-PET (Chromatin Interaction Analysis by Paired-End Tag Sequencing) (7) is a cutting-edge method that amalgamates chromatin immunoprecipitation (ChIP)-based enrichment, 3C, and high-throughput sequencing to pinpoint long-range chromatin contacts mediated by a transcription factor or a chromatin mark of interest. Concurrently, HiChIP and PLAC-seq integrate the benefits of ChIP and in situ Hi-C methodologies (8, 9). Compared to ChIA-PET, both HiChIP and PLAC-seq necessitate a reduced number of input cells and demonstrate heightened sensitivity, thereby presenting more efficient alternatives for chromatin interaction analysis.

In the intricate spatial organization of the three-dimensional (3D) genome, various functional proteins play a critical role in the formation and functional regulation of loop structures. The CCCTC-binding factor (CTCF) protein, through its interaction with DNA, forms a series of insulating loops comprised of CTCF-CTCF and CTCF-cohesin cross-binding interactions. These insulating loops contribute to maintaining the 3D structure of chromatin, modulating gene expression levels, and preserving genome stability (10). Structural Maintenance of Chromosomes (SMC) proteins are structural proteins that work in tandem with cohesins to intertwine different chromatin regions, forming chromatin loop structures (11). Histone H3 lysine 27 acetylation (H3K27ac) and Yin Yang 1 (YY1) proteins are associated with the activation of gene expression, functioning as structural regulatory factors for enhancer-promoter interaction, and promoting gene expression (12). Conversely, regions of the genome enriched with histone H3 lysine 27 trimethylation (H3K27me3) can act as silencers, suppressing gene expression through chromatin interactions (13). However, current targeted chromatin conformation capture experiments (such as ChIA-PET, HiChIP, and PLAC-seq) are limited in their capacity to capture loops mediated by limited number of target proteins and cell types, with data acquisition being resource-intensive, time-demanding, laborious, and subject to theoretical constraints.

Growing evidence suggests that DNA sequence and epigenomic features serve as reliable predictors of regulatory interactions and chromatin architecture. Utilizing these features, some computational methods have been established for predicting chromatin loops (14). Peakachu (15) is a Random Forest classification framework designed to predict chromatin loops from genome-wide contact maps. DeepYY1 (16) is a deep learning-based approach for predicting YY1 protein-mediated chromatin loops using genomic sequence features. ChINN (17) can predict CTCF- and RNA polymerase II-associated and Hi-C chromatin loops. RIPPLE (18), TargetFinder (19) and LoopPredictor (20) can detect cell type-specific enhancer-promoter loops by employing a wide range of features, including open chromatin measures, gene expression, transcription factors, and modified histones. Nonetheless, the necessity for diverse inputs makes it difficult to apply these methods to develop specific classifiers for additional cell types. In addition, none of current methods can predict various protein-mediated loops, this constraint poses a challenge for researchers attempting to unravel the complex interplay of various protein-mediated loops and their roles in shaping the 3D genome architecture and regulating gene expression across diverse cellular contexts. Consequently, advancing our understanding of the multitude of protein-mediated loop structures and their functional significance necessitates the development of novel experimental techniques and computational approaches that can concurrently capture and analyze multiple protein-mediated loops in various cell types.

In this study, we proposed a fusion neural network (FusNet) for predicting various protein-mediated loops. FusNet is composed of three layers: a feature extractor layer, predictor layer, and model fusion layer. The feature extractor layer employs a convolutional neural network to reduce dimensionality and extract information from genomic sequences. The predictor layer is constructed by integrating Light Gradient Boosting Machine, eXtreme Gradient Boosting and K-Nearest Neighbors models. The fusion layer utilizes a stacking strategy to fuse the prediction results from each basic model as new features for model training and predicts the final probability of loop formation. FusNet effectively predicts loops mediated by various proteins, the predictive results exhibit a high consistency with Hi-C data and are found to be linked to regulatory functions. Additionally, through association analysis with risk variants, FusNet has shown potential in uncovering disease-related mechanisms in complex disease research.

## Results

### DNase I hypersensitive and protein binding sites form protein-mediated loop anchors jointly

To investigate the performance of existing methods in predicting protein-mediated loops, we first conducted a comparative analysis of the loop prediction results of several existing approaches. Since the DeepYY1 method can only predict YY1 protein-mediated loops, we compared the YY1 protein-mediated loops generated by different methods and used available K562-YY1 HiChIP data as our standard. Initially, we performed an overlap analysis of loops predicted by the three methods with HiChIP loops. The overlap between loops obtained by the three methods and HiChIP loops was not satisfactory, with Peakachu achieving the highest overlap (less than 50%) among the three methods (**Fig. 1A**). Subsequently, we compared the overlap of predicted loop anchors with HiChIP loop anchors, and the results showed that ChINN achieved the highest anchor overlap (less than 70%) among the three methods (**Fig. 1B**). Next, we divided the predicted loops into bin intervals according to the anchors distances and compared the paired end tag (PET) proportions in different distance intervals with HiChIP data. The results demonstrated that the loop distance distributions predicted by the three methods were substantially different from the standard data (**Fig. 1C**). Since YY1 protein-mediated loops are closely related to enhancer regulation, we examined the loop distribution between the *SUFU* gene and nearby enhancer (21). The results indicated that none of the three methods predicted the association between the *SUFU* gene and the nearby enhancer (**Fig. 1D**). Similarly, near the gene *DDX19B*, none of the three prediction methods achieved satisfactory prediction results (*SI Appendix*, **Fig. S1A**).

**Figure 1.**
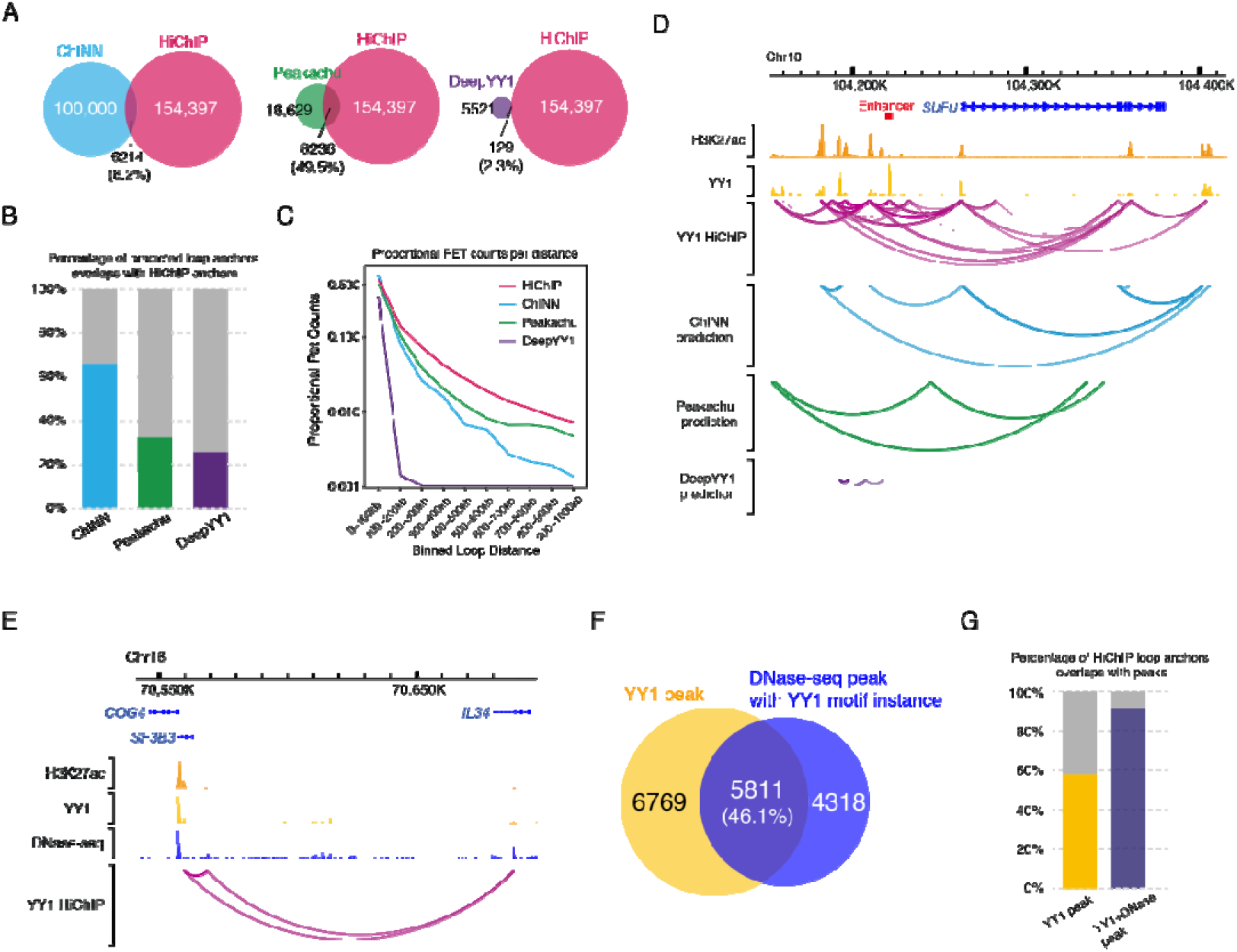
Combining DNase I hypersensitive sites and protein binding sites to identify protein-mediated loop anchors. **(A)** Overlap between loops obtained by the three predictive methods and HiChIP loops. **(B)** Comparison of loop anchors obtained by three predictive methods and derived from HiChIP data. **(C)** Distribution of PETs binned by loop distance in HiChIP dataset and the prediction of three predictive methods. **(D)** Tracks of HiChIP loops and the predicted loops of three predictive methods. **(E)** Tracks of H3K27ac, YY1 ChIP-seq peaks, DNase-seq peaks and HiChIP loops. The red squares indicate the anchor regions enriched with DNase-seq peaks. **(F)** Overlap between YY1 ChIP-seq peaks and DNase-seq peaks. **(G)** Percentage of HiChIP loop anchors overlaps with YY1 peaks and the merging results of YY1 peaks and DNase peaks.

To further understand the characteristics of protein-mediated loop anchors, we continued to analyze YY1 protein-mediated loops. By mapping H3K27ac, YY1 ChIP-seq peaks, DNase-seq peaks, and YY1 HiChIP loops to the genome, we found that the regions containing YY1 HiChIP loop anchors were not only related to YY1 ChIP-seq peaks but also to open chromatin regions. For example, the anchors located 20 kb downstream of the *SF3B3* gene and those within the *IL34* gene were enriched only in DNase-seq peaks (**Fig. 1E**). Similarly, near the gene *GSDME*, we observed an enrichment of DNase-seq peaks at loop anchors. (*SI Appendix*, **Fig. S1B**). Our overlap analysis of YY1 ChIP-seq peaks and DNase-seq peaks revealed that the number of DNase-seq peaks was significantly higher than that of YY1 ChIP-seq peaks, and the majority of YY1 ChIP-seq peaks (77.9%) were covered by DNase-seq peaks (**Fig. 1F**). Ideally, we believe that YY1 HiChIP loop anchors should have a high degree of overlap with YY1 ChIP-seq peaks; however, the analysis found that less than 60% of loop anchors overlapped with YY1 ChIP-seq peaks. We speculate that this is related to low levels of ChIP enrichment and restriction enzyme preferences in HiChIP experiments. When we merged and deduplicated the YY1 ChIP-seq peaks and DNase-seq peaks, the overlap with YY1 HiChIP loop anchors reached over 90% (**Fig. 1G**). Thus, combining YY1 ChIP-seq peaks and DNase-seq peaks can help identify protein-mediated loop anchors.

### Fusion Neural Network (FusNet) for predicting protein-mediated loops

FusNet is a deep learning model based on a fusion strategy, which takes genomic sequences as input, and outputs the predicted loops with a probability score. FusNet primarily comprises three components: the feature extractor layer, the predictor layer, and the model fusion layer (**Fig. 2**). The feature extractor layer employs a convolutional neural network (CNN) to capture the local and global features within the sequence through multiple convolution and pooling operations. By retaining the most prominent features while compressing their dimensions, the extractor reduces model complexity and enhances its generalization capabilities. Furthermore, the layer extracts meaningful information that aids in better understanding the structure and properties of the sequence data, ultimately contributing to more accurate predictions. The sequence features are captured by the feature extractor, and then combined with the distance between two anchors to generate new feature matrixes. Subsequently, the new features are input into the three models for independent predictions. The predictor layer is composed of three foundational models: Light Gradient Boosting Machine (LGB), eXtreme Gradient Boosting (XGB) and K-Nearest Neighbors (KNN). Each foundational model leverages its unique strengths to handle different data types and problems, thereby enhancing the overall predictive performance. The model fusion layer employs a stacking training approach, where the predictions of the three foundational models are fed into a meta-model, which integrates the outputs of the foundational models to obtain the final prediction outcome. This fusion strategy helps to reduce the errors and variances of individual models while enhancing the overall generalization capability. Through this approach, the FusNet model can more accurately predict the formation probability of loops within genomic sequences.

**Figure 2.**
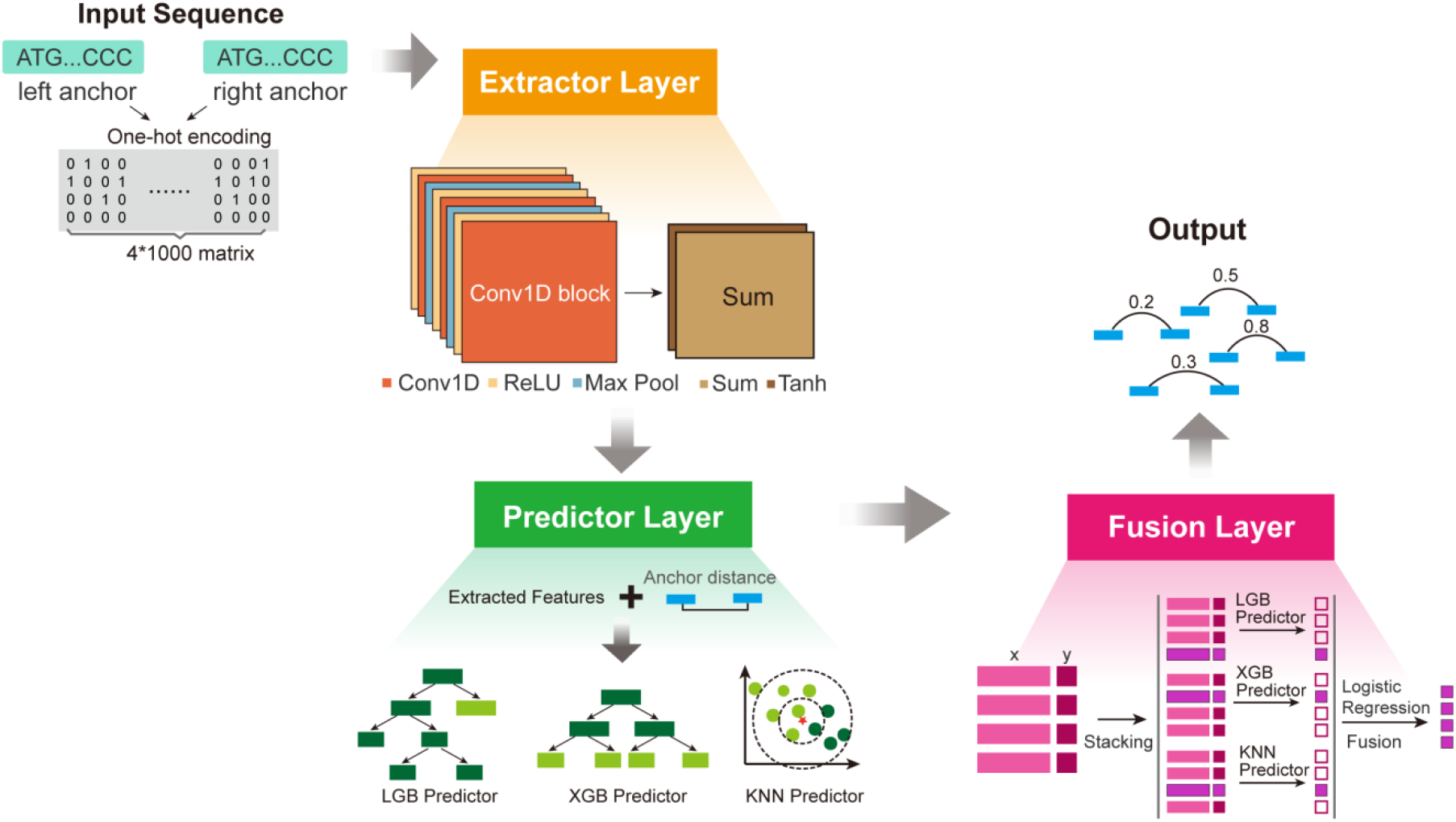
The schema of FusNet. FusNet primarily comprises three components: the feature extractor layer, the predictor layer, and the model fusion layer. FusNet takes sequence as input, and outputs the probability of predicted loops.

### Efficient feature capture with the feature extraction layer

By inputting data into the feature extractor layer and incorporating the extracted feature matrix into the fundamental model, we computed the average prediction results of the fundamental model and designated this process as FusNet-A. To analyze the feature extraction efficiency of FusNet’s feature extractor layer and study the importance of features, we conducted FusNet-A tests using sequences, anchor distances, and a combination of both features. To compare the features, we also constructed a random predictor, which assigns a random probability value between 0 and 1 to each pair of anchors and serves as a baseline for comparison. Comparative tests were conducted on eight different ChIP-seq datasets. The results showed that the model with both types of feature inputs achieved the highest auPRC values in all datasets, with auPRC values exceeding 0.8 in six datasets, except for Fgfr1 and ESR1 protein datasets. The performance of the model trained using sequence features alone was better than that of the model trained using anchor distances alone, but both significantly exceeded the randomly generated model (**Fig. 3A**). This indicates that, compared to anchor distances, sequence features contain more information and are more conducive to model learning.

**Figure 3.**
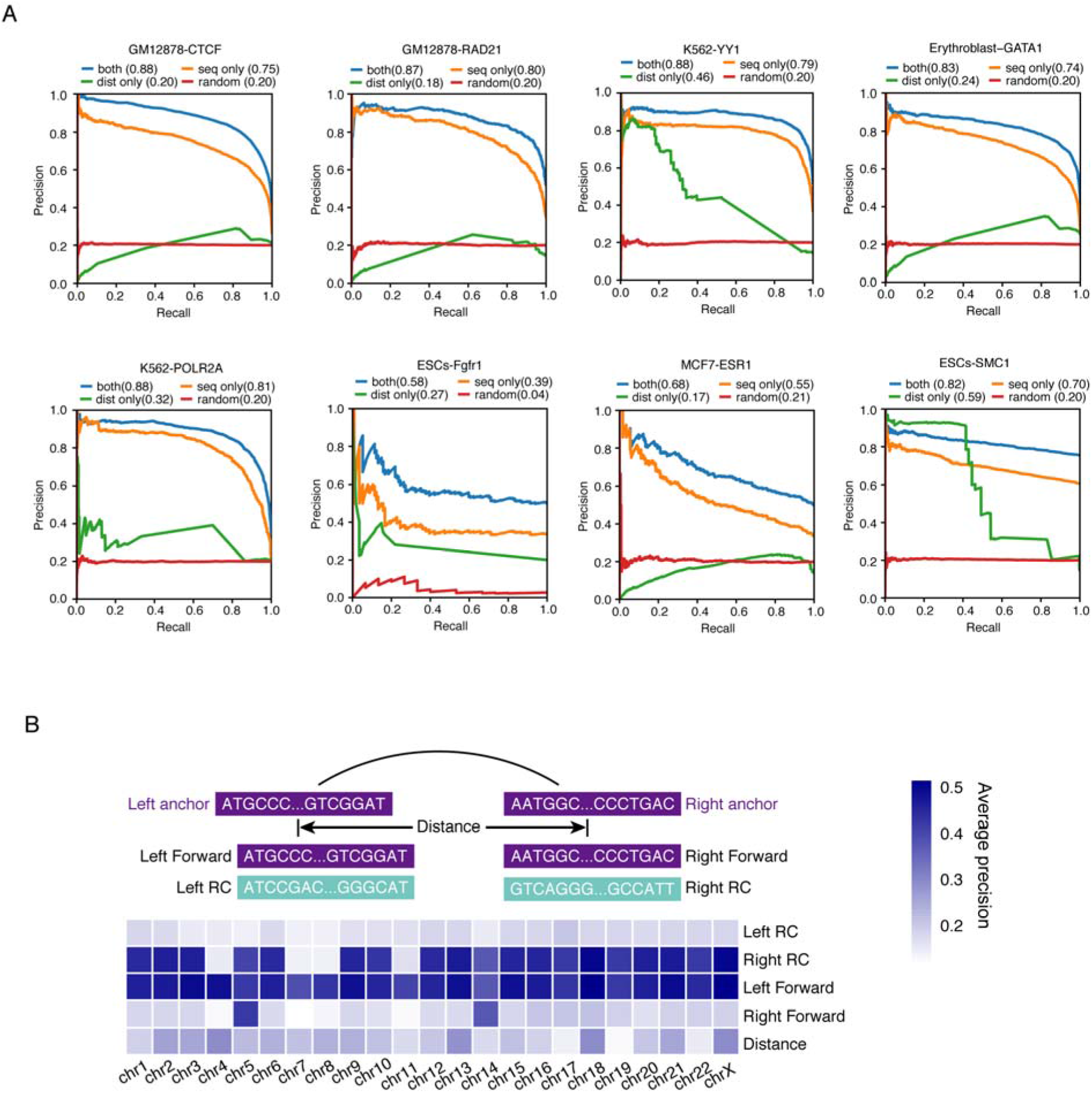
Investigation of feature importance. **(A)** The precision recall curve of FusNet-A trained with sequence, loop anchor distance and a combination of both data, the value of auPRC for each condition was showed inside bracket. **(B)** Assessing feature importance in various sequence extraction situations.

FusNet processes input genomic data into 4 sequences: the forward sequence of the left anchor, the reverse complement sequence of the left anchor, the forward sequence of the right anchor, and the reverse complement sequence of the right anchor. The reverse complement sequence is obtained by reversing the original sequence and replacing each base with its complementary base. The reverse complement of each sequence is unique. By applying permutation importance to the four sequence features on the 23 human chromosomes, we calculated the importance of different features in the predictions. Permutation importance is a model-agnostic method for measuring the contribution of a given feature to model performance. The results showed that the forward sequence of the left anchor and the reverse complement sequence of the right anchor achieved significantly higher importance among all features (**Fig. 3B**).

### Enhancing predictive performance with a fusion layer

To evaluate the effect of the fusion layer in FusNet on the model predictions, we named the complete process with the fusion layer as FusNet-F and compared it with FusNet-A which based on an average strategy. The results showed that except for the ESCs-Fgfr dataset, FusNet-F achieved a higher auPRC compared to FusNet-A on 7 different ChIP-seq datasets, indicating that the fusion layer could effectively improve the prediction performance of FusNet. (**Fig. 4A**). To investigate whether FusNet-F can learn useful information related to protein-mediated interactions, we applied the FusNet-F model trained on specific cell types to predict loops mediated by the same protein in different cell types. When we trained FusNet-F on the CTCF dataset corresponding to the HelaS3 cell type, we found that the prediction accuracy was unsatisfactory, with an auPRC of 0.64, which we attributed to the small amount of training data.

**Figure 4.**
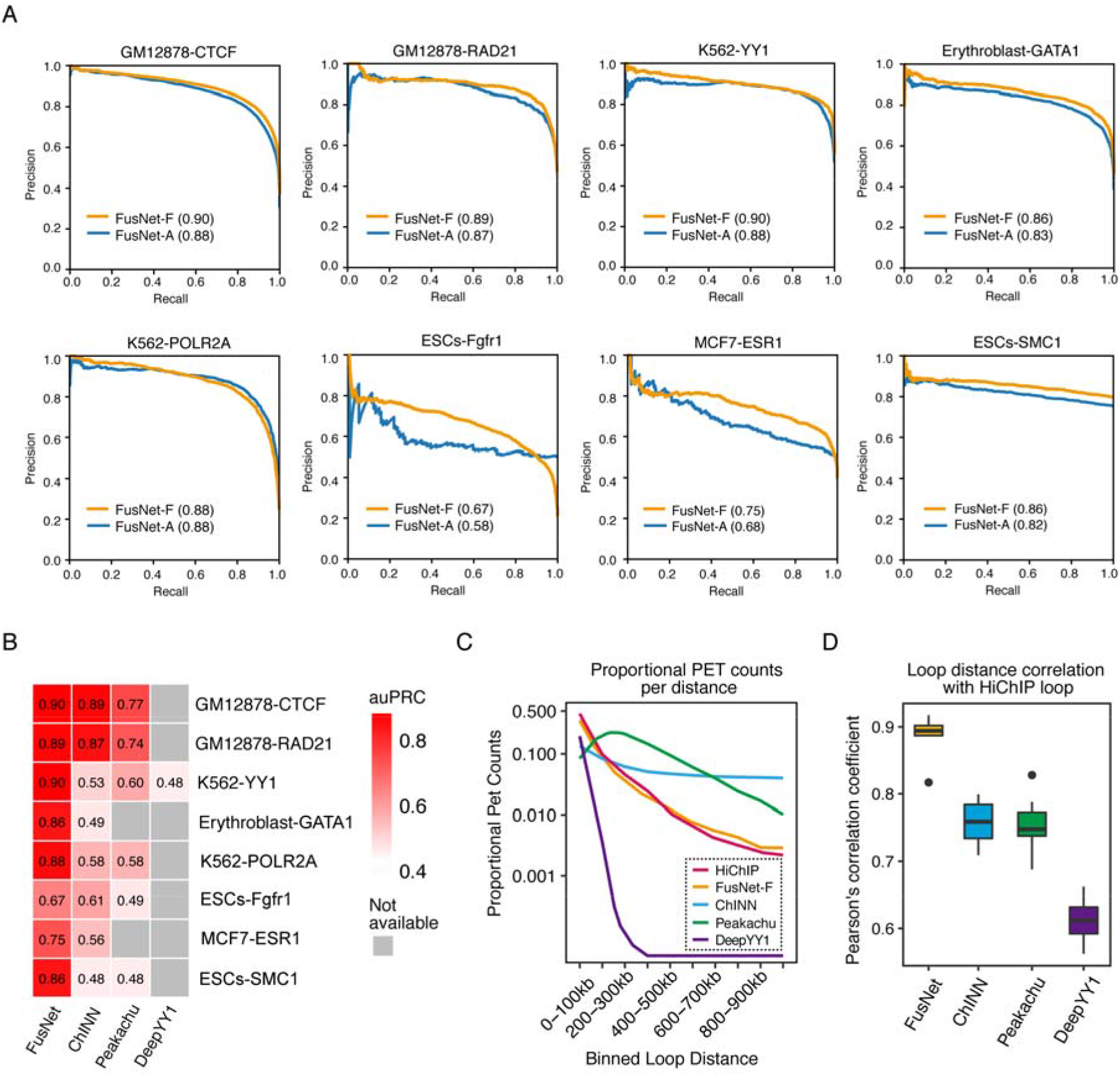
Improving predictive performance with fusion layer. **(A)** Evaluating prediction results of FusNet-A and FusNet-F on eight datasets, the value of auPRC were showed inside brackets. **(B)** Evaluating prediction results of FusNet-F and three other predictive methods on 8 datasets. the value of auPRC were showed inside heatmap, and the gray squares indicated that the data was not available. **(C)** Distribution of PETs binned by loop distance in HiChIP dataset and the prediction of four predictive methods. **(D)** Pearson’s correlation between the loop distance of four predictive methods and HiChIP loops.

However, when we trained FusNet-F on the CTCF dataset corresponding to the GM12878 cell type and performed cross-cell type prediction on CTCF-mediated loops in HelaS3, the prediction accuracy improved significantly. This indicated that the model fusion layer learned CTCF-mediated information from the GM12878 dataset and effectively applied it to cross-cell type prediction. Similarly, FusNet-F achieved significant improvements in cross-cell type loop prediction mediated by the RAD21 and POLR2A proteins (*SI Appendix*, **Fig. S2**).

To further evaluate the prediction performance of FusNet, we compared the results of FusNet with those of three other existing methods on eight datasets. The results showed that FusNet achieved higher auPRC values on all datasets (**Fig. 4B**). Additionally, we divided the loops predicted by different tools into bin sets based on the distance between anchors, and then calculated the distribution of loop numbers in different distance intervals. The results showed that for both single protein-mediated loop prediction and merged dataset prediction, the loop distribution of FusNet was closer to what we observed from HiChIP data (**Fig. 4C** and *SI Appendix*, **Fig. S3**). And we calculated the pearson’s correlation of loop distance between the predictive methods and HiChIP loops, FusNet showed highest correlation coefficient, indicating better prediction accuracy (**Fig. 4D**).

In addition, we also focused on what the FusNet model learned from the input data. We plotted the distribution of predicted results in each probability interval for FusNet trained on eight different transcription factor ChIP-seq datasets. Taking the best-performing GM12878-CTCF dataset as an example, most positive samples (56.8%) achieved a prediction probability greater than 0.5, while the model had better discriminatory power for negative samples. 95.8% of negative samples were assigned a prediction probability less than 0.1, and there were very few negative samples in other probability intervals. As for the falsely classified negative samples, there were almost none. Therefore, the CTCF model learned some important common features among negative samples from the input data. Through this, FusNet could predict most negative samples (*SI Appendix*, **Fig. S4**).

### Assessing FusNet predictions with Hi-C

To quantify how well each predicted loop set was supported by Hi-C data, we adopted Aggregate Peak Analysis (APA) which includes a measurement for P2LL (Peak to Lower Left) (22) to calculate the aggregate enrichment of a set of putative peaks within a contact matrix. A strong dark pixel at the center of an APA plot signifies the specific enrichment of Hi-C contacts concerning loop calls relative to their local background. The enrichment is further quantified by APA P2LL scores, the ratio of the central pixel to the mean of the pixels in the lower left corner. The results demonstrated that FusNet achieved high APA P2LL scores (>2.5) for predictions across all eight ChIP-seq datasets (**Fig. 5A**).

**Figure 5.**
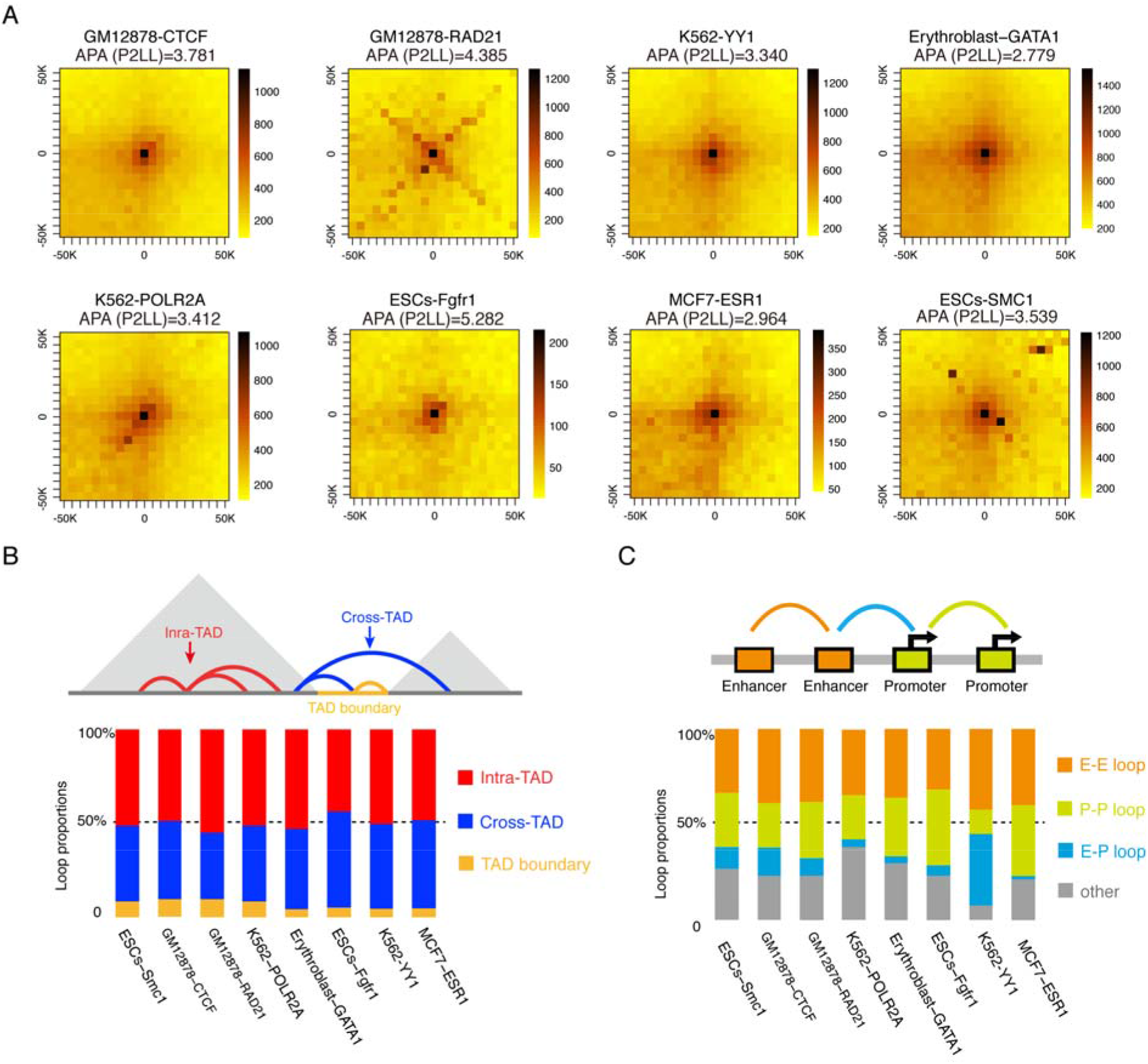
Evaluating FusNet predictions. **(A)** APA plot with P2LL measurement on eight datasets. **(B)** Percentage of loops in different TAD localization. The curves in the diagram indicated three TAD localization situations. **(C)** Functionality annotation of predicted loops.

Loops and topologically associating domains (TADs) are essential hierarchical structures in the three-dimensional chromatin organization, playing crucial roles in gene regulation and genome stability. In recent years, numerous studies have shown that although TAD structures exhibit insulating properties that concentrate chromatin interactions within TADs, many loops still span TADs for distal gene regulation. Such loop structures are closely related to transcriptional regulation (23). Additionally, loops are enriched within TAD boundaries, which are typically rich in CTCF and cohesin protein complexes and exhibit a more active open chromatin state. These boundaries are conserved across different species, and the maintenance of TAD boundaries contributes to the stability and specificity of gene expression by restricting chromatin interactions within their respective regions, preventing interactions between different TADs from affecting gene expression regulation (24). By mapping the loops predicted by FusNet on eight different ChIP-seq datasets to the published TAD structures, we obtained the proportion of loops in different TAD regions, providing further insight into the structural properties of the prediction results. The results showed that in the eight datasets, most loops were located within TADs (Intra-TAD), followed by those spanning TAD regions (Cross-TAD), and only a small number of loops were situated at TAD boundaries (**Fig. 5B**). This demonstrates that the prediction results of FusNet are in general agreement with current research findings.

Interactions between enhancers and promoters play a vital role in the regulation of gene expression. To analyze the prediction results from a functional perspective, we first collected enhancer and promoter regions for relevant cell lines from ENCODE. Next, we mapped the predicted loops to the enhancer and promoter sets of corresponding cell line, annotating loop types accordingly. The annotation types are classified as Enhancer-Enhancer (E-E), Promoter-Promoter (P-P), Enhancer-Promoter (E-P), and other. Any loop with neither anchor belonging to an enhancer or promoter is classified as “other”. The loop type annotation results show that the K562-YY1 loop predictions are primarily E-E and E-P types, with the fewest “other” type loops. This is consistent with the functional characteristics of the YY1 protein, which activates gene expression. In addition, proteins such as Smc1, CTCF, RAD21, and POLR2A form cohesin protein complexes and are associated with loop structure formation. In the loop predictions corresponding to these proteins, the “other” type occupies a larger proportion (**Fig. 5C**).

### Applying FusNet to uncover the role of disease-associated risk variants

Single nucleotide polymorphisms (SNPs) are often located in distal intergenic regions but may impact gene expression by interacting with expressed gene targets through chromatin loops (25). Therefore, long-range chromosomal contacts may explain how SNPs within distal regulatory elements contribute to the risk of certain diseases. In this study, we collected 516,246 SNPs from 18 complex diseases and traits in the GWAS Catalog (26) and mapped them onto merged predicted loops from the FusNet model. We counted the loops for 50 upstream and downstream anchor fragments of each SNP and found that loop numbers were significantly higher near SNP locations than in other regions (**Fig. 6A**). Similar results were obtained in individual dataset predictions from FusNet (*SI Appendix*, **Fig. S5A**). We also classified all SNPs into non-interacting and interacting GWAS SNPs based on whether they were covered by a loop or not and calculated the ratio of median p-values for 18 complex diseases and traits. The results showed that SNPs related to metabolic disorders and hematological measurements were more likely to be located in FusNet predicted loop anchors than other SNPs (**Fig. 6B**). Similar results were also observed in individual data set predictions (*SI Appendix*, **Fig. S5B**).

**Figure 6.**
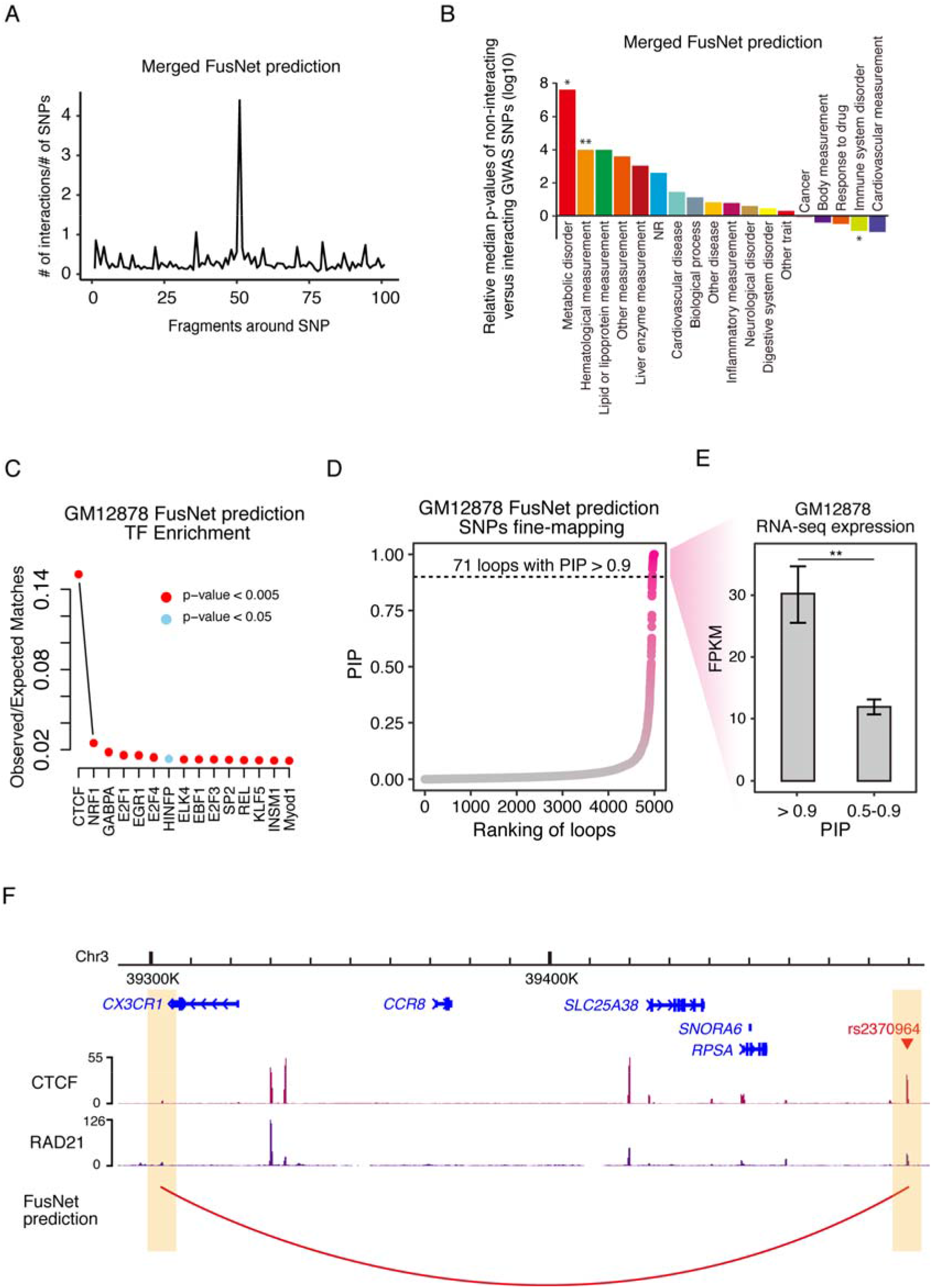
Uncovering the regulatory role of FusNet predicted loops. **(A)** Counting of merged FusNet loops near SNPs positions. **(B)** Statistical significance of interacting and non-interacting GWAS SNP fragments. Difference between interacting and non-interacting GWAS SNP fragments was calculated with T test, * p-value < 0.05, ** p-value < 0.01. **(C)** Transcription factors enrichment of FusNet prediction associated SNPs on GM12878 dataset. **(D)** SNPs fine-mapping of FusNet predicted loops on GM12878 dataset. **(E)** RNA-seq expression of SNPs target genes within different range of PIP scores. Difference between two conditions was calculated with T test, ** p-value < 0.01. **(F)** Example of FusNet predicted loops. The yellow squares indicated the anchors regions of observed loop.

To further analyze the SNPs located within loops predicted by FusNet, we extracted SNPs from predicted loops in the GM12878 cell line and performed transcription factor enrichment analysis. The results showed that the enrichment level of CTCF was significantly higher than other transcription factors (**Fig. 6C**). Additionally, we mapped predicted loops in GM12878 cells to eQTL data in blood tissue and extracted overlapping loops. These loops were sorted according to the posterior including probabilities (PIP) score, and analysis showed that 71 loops in GM12878 cells had a high (>0.9) PIP score, more than that of K562 and MCF7 (**Fig. 6D** and *SI Appendix*, **Fig. S6A-S6B**). Other cell lines were not analyzed because of the lack of corresponding eQTL data. Subsequently, we used RNA-seq data for expression analysis, and the results showed that the expression level of target genes corresponding to SNPs with a PIP score greater than 0.9 was significantly higher than that of target genes corresponding to SNPs in the range of 0.5-0.9 (**Fig. 6E** and *SI Appendix*, **Fig. S6C-S6D**). Finally, we discovered that the risk variant rs2370964 on chromosome 3 interacts with the promoter region of the distant gene *CX3CR1* (**Fig. 6F**).

Consistent with previous reports (27), rs2370964 is associated with the expression level of *CX3CR1*, and this SNP may contribute to the risk of amyotrophic lateral sclerosis (ALS). These findings suggest that the predictive results of FusNet may provide new insights into pathogenesis, and the proposed research methodology can be applied to other complex diseases as well.

## Discussion

In the three-dimensional (3D) genome, the intricate spatial organization of loop structures is regulated by various functional proteins. However, current chromatin conformation capture experiments are limited in their ability to capture loops mediated by a limited number of target proteins and cell types. Obtaining data is resource-intensive, time-consuming, and subject to theoretical constraints, and no current methods can predict various protein-mediated loops. This presents a challenge for researchers attempting to unravel the complex interplay of protein-mediated loops and their roles in shaping the 3D genome architecture and regulating gene expression across diverse cellular contexts. To advance our understanding of the multitude of protein-mediated loop structures and their functional significance, it is necessary to develop novel experimental techniques and computational approaches that can concurrently capture and analyze multiple protein-mediated loops in various cell types.

In this study, we propose FusNet, a model fusion-based method for predicting various protein-mediated loops. FusNet consists of a feature extractor, predictor, and model fusion layer. It takes the genome sequence as input, determines anchor positions through open chromatin and ChIP-seq data, and uses one-hot encoding to transform the genome sequence into a matrix. The feature extractor uses a convolutional neural network to reduce dimensionality and extract information from the feature matrix, and uses a weighted sum layer to obtain the summary scores of each kernel in the last convolutional layer. This allows each anchor, regardless of its size, to be represented by the same number of features. The predictor layer is composed of three basic models: Light Gradient Boosting Machine, eXtreme Gradient Boosting and K-Nearest Neighbors. The fusion layer adopts a stacking strategy, where the meta-model is employed to combine the predictions from all fundamental models and predict the probabilities of loop formation.

To investigate the importance of different sequence features, we calculated the permutation importance, which showed that the forward sequence of the left anchor and the reverse complementary sequence of the right anchor achieved the highest importance scores among all features. It is noteworthy that these two sequences converge in direction, similar to the motif enrichment direction of CTCF and other structural proteins reported in previous literature on loop anchor points (28). Additionally, the distance between the anchor points also has some helpful effect on improving the prediction performance, although not significantly so.

To evaluate the effectiveness of the fusion layer based on stacking strategy for prediction, we replaced the fusion layer of FusNet-F with the average method, and constructed the FusNet-A model. The comparison between FusNet-F and FusNet-A showed that FusNet-F achieved better prediction results in all datasets, and it could effectively predict cross cell type protein-mediated loops. This suggests that the fusion layer based on the stacking strategy is helpful for improving the prediction performance of the model.

To further analyze the role of chromatin interactions in identifying potential target genes of pathogenic single nucleotide polymorphisms (SNPs), we mapped and quantitatively analyzed the genomic coordinates of SNPs with FusNet predicted loop anchors. The results showed that in all eight datasets, the number of loops near SNPs exhibited an enrichment, indicating the aggregation of SNPs in the vicinity of loop anchors. This may be because the formation of chromatin Loop structures often implies the presence of functional elements such as enhancers, promoters, and insulators, which play important roles in gene regulation. Genetic variations (such as SNPs) surrounding these functional elements may affect gene expression and regulation.

Besides, loop anchors may be located in highly conserved genomic regions that have been relatively stable during evolution across different species due to their important biological functions. In these conserved regions, genetic variations (such as SNPs) may experience stronger selective pressures, leading to the aggregation of functional SNPs.

## Materials and Methods

### Processing of HiChIP and ChIA-PET data

For the HiChIP datasets, we used HiC-Pro (29) to preprocess the raw reads, then retained the high-quality alignments with MAPQ score of 30. The valid pairs were input to hichipper with the loop calling mode “Each+Self”, and the output loops were filtered with at least 5 paired end tags, and p-value < 0.05. For ChIA-PET datasets, we used ChIAPoP (30) to preprocess the raw reads and the minimum MAPQ score was set to 30. In the loop calling step, the minimum counts was set to 3 and p-value threshold was to 0.05 to filter the final loops.

### One-hot Encoding for FusNet Model Inputs

The FusNet model takes the sequence of anchor pairs as input, firstly, the sequence is converted to a 4*1000 matrix through one-hot encoding, which is a method for numerical representation of categorical variables. The basic idea of one-hot encoding is to transform each categorical value into a binary vector of length equal to the number of categories, with only one element being 1 and the others being 0. In this study, the four types of nucleotides (A, T, G, C) are represented by four-dimensional vectors, where nucleotide A is represented by [1, 0, 0, 0]^T^, G by [0, 1, 0, 0]^T^, C by [0, 0, 1, 0]^T^, and T by [0, 0, 0, 1]^T^. Each column of the matrix represents a nucleotide, while 1000 denotes the sequence interval consisting of 1000 nucleotides. If a sequence exceeds 1000 nucleotides, it is divided into multiple adjacent intervals to ensure that the number of nucleotides in each interval does not exceed 1000. For unknown nucleotides, such as NaN, they are assumed to have an equal probability of occurrence among the four nucleotide types, and therefore are represented as [0.25, 0.25, 0.25, 0.25]^T^.

### Feature extractor layer

The feature extraction layer is constructed by Convolutional Neural Networks (CNN), where the convolutional layer consists of multiple kernels. Each kernel performs cross-correlation independently with the input to measure the local similarity between the input and the kernel. In the FusNet model, a weighted sum layer is used after the last convolutional layer to obtain the summary score of each kernel. The weighted sum layer learns the weight distribution of each channel in the input independently and calculates the weighted sum of each channel along the spatial dimension. Intuitively, the weighted sum layer learns to weight the detected features in different spatial regions. For example, it may assign higher weights to the features detected near the center of the input sequence while giving negative weights to the features detected in the surrounding areas for punishment. A non-linear activation function such as tanh or sigmoid is applied after the weighted sum layer. The size of loop anchors varies greatly from thousands to tens of thousands of base pairs. Therefore, the network needs to adapt to different anchor sizes. For each anchor, it is divided into sub-intervals of 1000 bp with a 500 bp overlap between consecutive sub-intervals. The sequence of each sub-interval is independently fed into the convolutional layer. After the non-linear function following the weighted sum layer, the maximum value of each kernel in all sub-intervals of the anchor is taken as the feature value of the anchor. Therefore, each anchor, regardless of its size, is represented by the same number of features.

Here, the anchor can be represented by a matrix *X* with the shape of *N* × *L* × *C*, where *N* is the number of sub-intervals, *L* is the length of each sub-interval, and *C* is the number of channels (or nucleotides). For each kernel in the convolutional layer, let *S* be the width of the kernel, *W* ^*k*^ be the weight matrix of size *S* × *C*, and *b*^*k*^ be the bias. The convolution operation is as following,

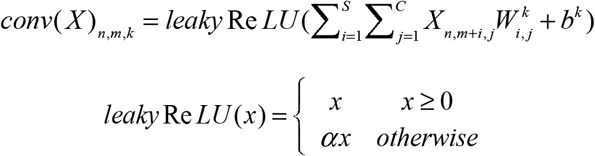

For the max pooling, let *S* be the size of the pool and *T* be the stride. Then, for an input matrix *X* of shape *N* × *L*× *C*, the computation of the maximum value in the local region is as following,

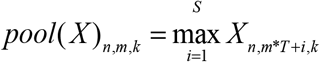

The output of the weighted sum layer is of shape *N* × *C* . For the weighted sum layer, let *W*^*ws*^ be the weight matrix of shape *L* × *C*, the calculation of the weighted sum is as following,

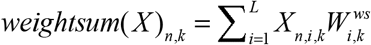

### Predictor layer

In the predictor layer, LGB (31), XGB (32), and KNN (33) are combined to form the base models. These models are trained on new features that are a combination of the features extracted by the feature extractor and the distance between anchors. The training process of the model consists of two steps. In the first step, the feature extractor is trained using an initial dataset. Positive anchor pairs are sampled, and similar negative samples are generated according to the sampling results. In this step, a two-layer fully connected neural network is constructed as a classifier. After receiving the features extracted by the feature extractor, the classifier will output the predicted probability of forming a loop. During the training of the feature extractor, multiple rounds of training are conducted on a batch of data. The training loss of the current batch is calculated, and the Adam optimizer is used for backpropagation to update the model’s parameters. After training the current epoch, a prediction is made using the validation set to obtain the validation loss. If the validation loss of the current epoch is lower than the best historical validation loss, the model and classifier are saved, and the best validation loss is updated. If the validation loss has not decreased further in the last three rounds, the training is stopped. In the second step, more negative samples are added to the initial dataset to obtain an augmented dataset. The best-performing feature extractor obtained in the first step is used to extract features, and the three base models of the predictor layer are trained on the expanded dataset. The outputs of three base models are independent of each other, and the probability of forming a loop for two anchors is within the range of 0 to 1.

### Fusion Layer

Fusion layer combines the predictions of base models to improve overall performance. Here we use stacking to implement the fusion process, which takes the predictions of multiple different basic models as inputs and trains a new meta-model to perform fusion and obtain better prediction performance. Stacking has many advantages, such as reducing overfitting by combining the predictions of multiple models and leveraging the strengths of each model to improve the final prediction accuracy. Additionally, stacking can improve the robustness of the final prediction by reducing the variance of each model.

The predictor layer trains three base models, denoted as *L*_*XGBoost*_, *L*_*SVC*_, and *L*_*KNN*_, and obtains their respective predictions for an input sample *X*, denoted as *y*_1_ = *L*_*XGBoost*_ (*X*), *y*_2_ = *L*_*SVC*_ (*X*), and *y*_3_ = *L*_*KNN*_ (*X*). These predictions are combined into a new feature vector *Y*_*base*_ = [*y*_1_, *y*_2_, *y*_3_], which is then used as input for the meta-model, which maps the output of the linear model to the (0,1) interval using the Sigmoid function. The resulting output value is used as the probability of forming a loop, which is calculated as following,

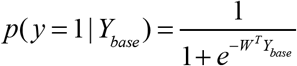

### Calculation of Permutation Importance

To calculate the permutation importance, we randomly shuffle the values of a feature and then re-calculate the model’s performance to measure the importance of that feature. If a feature has a large impact on the model’s performance, shuffling its values will significantly decrease the model’s performance. The steps of the permutation importance are as follows: first, train a model and evaluate its performance metric, such as accuracy; for each feature in the model, shuffle its values by randomly swapping the feature’s values with values from other samples and re-evaluate the model’s performance metric, calculate the degree to which the model’s performance decreases, and use that as the feature’s importance score. Finally, rank all the feature’s importance scores to determine their relative contribution to the model’s performance.

### Mapping loops to TADs

To map predicted loops to TAD and obtain the proportion of loops in different TAD regions, we first downloaded the corresponding Hi-C dataset for the cell line from the 4DN data portal (34), and then identified insulation domains using FAN-C (35) call with a window size of 1 Mb. Here, we overlapped the anchors at both ends of the loop with insulation domains. If both anchors were in the same domain, we defined it as an intra-TAD loop. If both anchors were in the same boundary region, we defined it as a TAD boundary. Otherwise, if the anchors crossed at least one domain region, we defined it as a cross-TAD loop.

## Availability

The source code of 3DFunc is available at GitHub repository: https://github.com/CSUBioGroup/FusNet

## Acknowledgments

We are grateful to the High-Performance Computing Center of Central South University for partial support of this work. This work was supported by grants from the National Natural Science Foundation of China under Grants (No. 62225209) [M.L.], Hunan Provincial Science and Technology Program (2019CB1007) [M.L.], the 111 Project (No. B18059) [M.L.].

## References

1. J. O. J. Davies, A. M. Oudelaar, D. R. Higgs, J. R. Hughes, How best to identify chromosomal interactions: a comparison of approaches. Nat Methods 14, 125–134 (2017).

2. M. Bulger, M. Groudine, Functional and Mechanistic Diversity of Distal Transcription Enhancers. Cell 144, 825 (2011).

3. J. Kooren, M. Simonis, W. de Laat, An evaluation of 3C-based methods to capture DNA interactions. Nat Methods 4, 895 (2007).

4. H. Hagège, et al., Quantitative analysis of chromosome conformation capture assays (3C-qPCR). Nat Protoc 2, nprot.2007.243 (2007).

5. A. Denker, W. de Laat, The second decade of 3C technologies: detailed insights into nuclear organization. Gene Dev 30, 1357–1382 (2016).

6. H. J. G. van de Werken, et al., Robust 4C-seq data analysis to screen for regulatory DNA interactions. Nat Methods 9, 969 (2012).

7. M. J. Fullwood, et al., An oestrogen-receptor-α-bound human chromatin interactome. Nature 462, 58 (2009).

8. M. R. Mumbach, et al., HiChIP: efficient and sensitive analysis of protein-directed genome architecture. Nat Methods 13, 919–922 (2016).

9. R. Fang, et al., Mapping of long-range chromatin interactions by proximity ligation-assisted ChIP-seq. Cell Res 26, 1345 (2016).

10. E. D. Rubio, et al., CTCF physically links cohesin to chromatin. Proc National Acad Sci 105, 8309–8314 (2008).

11. T. L. Higashi, F. Uhlmann, SMC complexes: Lifting the lid on loop extrusion. Curr Opin Cell Biol 74, 13–22 (2022).

12. A. S. Weintraub, et al., YY1 Is a Structural Regulator of Enhancer-Promoter Loops. Cell 171, 1573-1588.e28 (2017).

13. Y. Cai, et al., H3K27me3-rich genomic regions can function as silencers to repress gene expression via chromatin interactions. Nat Commun 12, 719 (2021).

14. H. Tao, et al., Computational methods for the prediction of chromatin interaction and organization using sequence and epigenomic profiles. Brief Bioinform, bbaa405 (2021).

15. T. J. Salameh, et al., A supervised learning framework for chromatin loop detection in genome-wide contact maps. Nat Commun 11, 3428 (2020).

16. F.-Y. Dao, et al., DeepYY1: a deep learning approach to identify YY1–mediated chromatin loops. Brief Bioinform 22 (2020).

17. F. Cao, et al., Chromatin interaction neural network (ChINN): a machine learning-based method for predicting chromatin interactions from DNA sequences. Genome Biol 22, 226 (2021).

18. S. Roy, et al., A predictive modeling approach for cell line-specific long-range regulatory interactions. Nucleic Acids Res 43, 8694–8712 (2015).

19. S. Whalen, R. M. Truty, K. S. Pollard, Enhancer–promoter interactions are encoded by complex genomic signatures on looping chromatin. Nat Genet 48, ng.3539 (2016).

20. L. Tang, et al., Predicting unrecognized enhancer-mediated genome topology by an ensemble machine learning model. Genome Res 30, 1835–1845 (2020).

21. J. Y. Ko, S. Oh, K. H. Yoo, Functional Enhancers As Master Regulators of Tissue-Specific Gene Regulation and Cancer Development. Mol Cells 40, 169–177 (2017).

22. N. C. Durand, et al., Juicer Provides a One-Click System for Analyzing Loop-Resolution Hi-C Experiments. Cell Syst 3, 95–98 (2016).

23. T.-H. S. Hsieh, et al., Enhancer–promoter interactions and transcription are largely maintained upon acute loss of CTCF, cohesin, WAPL or YY1. Nat Genet 54, 1919–1932 (2022).

24. E. McArthur, J. A. Capra, Topologically associating domain boundaries that are stable across diverse cell types are evolutionarily constrained and enriched for heritability. Am J Hum Genetics 108, 269–283 (2021).

25. M. Behrends, O. Engmann, Loop Interrupted: Dysfunctional Chromatin Relations in Neurological Diseases. Frontiers Genetics 12, 732033 (2021).

26. A. Buniello, et al., The NHGRI-EBI GWAS Catalog of published genome-wide association studies, targeted arrays and summary statistics 2019. Nucleic Acids Res 47, D1005–D1012 (2019).

27. A. Yousefian-Jazi, et al., Functional fine-mapping of noncoding risk variants in amyotrophic lateral sclerosis utilizing convolutional neural network. Sci Rep-uk 10, 12872 (2020).

28. Z. Tang, et al., CTCF-Mediated Human 3D Genome Architecture Reveals Chromatin Topology for Transcription. Cell 163, 1611–1627 (2015).

29. N. Servant, et al., HiC-Pro: an optimized and flexible pipeline for Hi-C data processing. Genome Biol 16, 259 (2015).

30. W. Huang, M. Medvedovic, J. Zhang, L. Niu, ChIAPoP: a new tool for ChIA-PET data analysis. Nucleic Acids Res 47, gkz062.(2019).

31. G. Ke, et al., LightGBM: A Highly Efficient Gradient Boosting Decision Tree in Advances in Neural Information Processing Systems, (Curran Associates, Inc.).

32. T. Chen, C. Guestrin, XGBoost: A Scalable Tree Boosting System. CoRR abs/1603.02754 (2016).

33. T. Cover, P. Hart, Nearest neighbor pattern classification. Ieee T Inform Theory 13, 21–27 (1967).

34. S. B. Reiff, et al., The 4D Nucleome Data Portal as a resource for searching and visualizing curated nucleomics data. Nat Commun 13, 2365 (2022).

35. K. Kruse, C. B. Hug, J. M. Vaquerizas, FAN-C: a feature-rich framework for the analysis and visualisation of chromosome conformation capture data. Genome Biol 21, 303 (2020).

